# Adaptive consequences of transgenerational inheritance of a predatory mouth-form trait in nematodes

**DOI:** 10.1101/2025.10.05.680509

**Authors:** Shiela Pearl Quiobe, Ralf J. Sommer

## Abstract

A growing body of evidence suggests that environmentally-induced changes in phenotypes can cross the generation barrier and associated molecular mechanisms are being increasingly identified. Such transgenerational epigenetic inheritance (TEI) is repeatedly considered to be adaptive although experimental support for this claim is often missing. We study the adaptive consequences of TEI in the facultative predatory nematode *Pristionchus pacificus*, which can form two alternative mouth forms dependent on environmental conditions. The choice of feeding structure in *P. pacificus* can be manipulated through dietary switching and food reversal. Recent studies showed that certain diets cause the induction of the predatory mouth form, which can result in TEI after food reversal. This memory requires the ubiquitin ligase EBAX-1/ZSWIM8 and the destabilization of clustered microRNAs. To study the adaptive potential of transgenerational memory of the predatory morph, we measured brood size as proxy for fitness. We compared naïve worms, with those under dietary induction and food reversal, and found a fitness advantage for animals exhibiting TEI of the predatory morph. Specifically, after food reversal worms had higher lifetime reproductive success for up to three generations when compared to diet-induced or naïve worms. These results support the adaptive significance of non-genetic inheritance of the predatory trait.

## Introduction

The environment is long known to influence organismal phenotypes and affect physiological, morphological and behavioral traits (West-Eberhard, 2003). Such plasticity is normally not transmitted to the next generation; thus, representing an intragenerational phenomenon. However, growing evidence reveals examples of both intergenerational and transgenerational inheritance, the latter referring to heritable effects that last for at least three generations beyond the original environmental exposure (for reviews see (Heard & Martienssen, 2014; Houri-zeevi & Rechavi, 2017; Baugh & Day, 2020; Sengupta, Kaletsky & Murphy, 2023)). This non-genetic form of inheritance in the absence of the original stimulus is alluded to as ‘transgenerational epigenetic inheritance’ (TEI), and is extremely widespread with examples from plants, nematodes, flies and mice (Chen & Rechavi, 2022). The molecular machinery associated with TEI of acquired characters is beginning to be identified, which involves a variety of epigenetic mechanisms, i.e. DNA methylation, histone modifications and small RNA signaling (Frolows & Ashe, 2021; Sengupta, Kaletsky & Murphy, 2023). In contrast, the biological and evolutionary significance of TEI is still contentious (Santilli & Boskovic, 2023). Remaining skepticism often stems from the fact that TEI is considered adaptive without proper experimental support for such claims. Therefore, a recent review commentary established three necessary criteria to document adaptive multigenerational plasticity and TEI (Baugh & Day, 2020). First, the demonstration that an environmental effect is transmitted across at least three generations. Second, an explanation for the mechanistic basis of the transgenerational phenomenon. And finally, experimental evidence that such multigenerational plasticity improves fitness in the descendants (Baugh & Day, 2020). Indeed, multiple studies in the last decade have found ample of evidence for criteria 1 and 2, i.e. the demonstration of the transmission of environmental information across multiple generations and associated molecular mechanisms. In contrast, direct proof for improved descendant fitness and thus, clear demonstration of adaptive multigenerational plasticity remains scarce. Here, we study the potential adaptive consequences of TEI of a predatory mouth-form trait in the self-fertilizing nematode *Pristionchus pacificus*.

Androdioecious hermaphroditic nematodes, such as *Caenorhabditis elegans* and *P. pacificus*, have several advantages for experimental research, including i) a short generation time (3-4 days at 20 °C), ii) simple husbandry on monoxenic bacterial diets (i.e. *Escherichia coli* or environmental-derived bacteria), and iii) propagation of isogenic cultures with genetically identical animals due to self-fertilization of hermaphrodites (Brenner, 1974; Schroeder, 2021). With its mouth-form plasticity and the ability to form two alternative feeding structures in response to environmental variation, *P. pacificus* provides a natural readout with ecological relevance that is deeply anchored in the natural history of the organism (for review see Sommer, 2025). During postembryonic development, genetically identical worms have to decide to either form the stenostomatous (St) morph with a narrow mouth and single dorsal tooth or the eurystomatous (Eu) form with a wide mouth and two teeth (Bento et al., 2010) (Figure 1A). Mouth-form plasticity has behavioral implications as only Eu animals are potential predators of other nematodes (Figure 1B) (Ishita et al., 2021; Quach & Chalasani, 2022; Eren et al., 2024). This natural example of developmental plasticity has emerged as one of the study systems in plasticity research because it can be characterized as a bi-stable developmental switch (Ragsdale et al., 2013; Kalirad & Sommer, 2024). This switch is inherently stochastic: even isogenic animals under fixed environmental conditions form both mouth forms in a genotype-specific ratio, i.e. the *P. pacificus* wild type strain PS312 from California is 90%:10% Eu:St under normal culture conditions, whereas other wild isolates show different ratios (Ragsdale et al., 2013; Dardiry et al., 2023). These characteristics allow mouth-form plasticity to be manipulated by forward and reverse genetic analyses and the associated gene regulatory network (GRN) is studied in great detail (Casasa, et al., 2023; Levis & Ragsdale, 2023; Reich et al., 2025). Similarly, epigenetic mechanisms regulating *P. pacificus* mouth-form plasticity are being identified (Werner et al., 2023; Brown et al., 2024).

**Figure 1.**
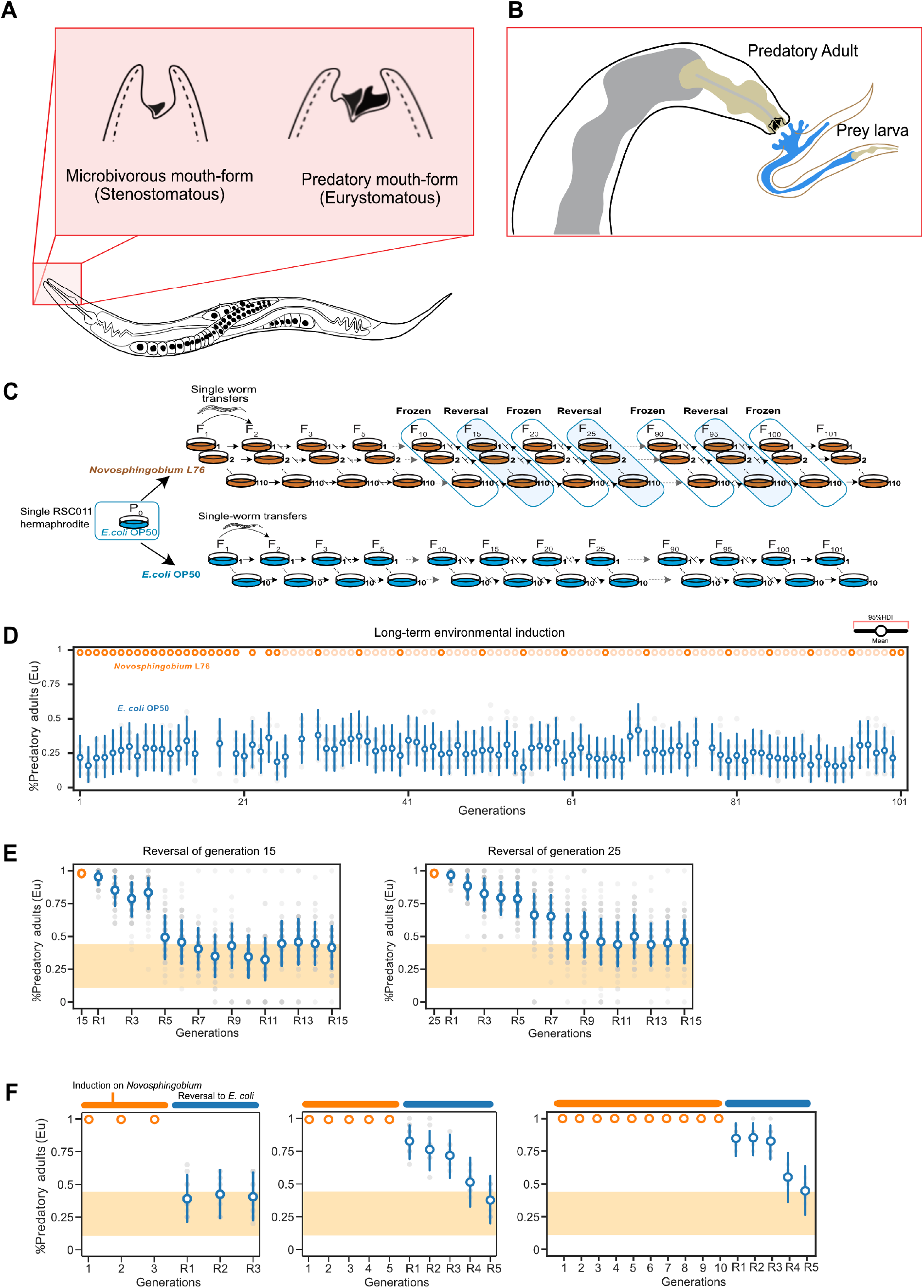
Overview about *P. pacificus* predation and long-term environmental induction experiment. **(A)** Schematic of the mouth dimorphism in *P. pacificus*. Stenostomatous animals have a single tooth and narrow mouth resulting in strict microbivorous feeding. In contrast, eurystomatous animals have two teeth and a broader mouth that allows both, microbivorous and predatory feeding. (**B)** Schematic of predation by a *P. pacificus* adult against a prey nematode of another species. (**C)** Experimental design of the original long-term environmental induction experiment. One hundred ten lines derived from the same *P. pacificus* RSC011 mother were propagated for 101 generations by single worm transfer on *Novosphingobium* L76 bacteria. Ten control lines were cultured on *E. coli* OP50 for the same period also by single worm transfer. Lines were frozen every 10 generations and were periodically reversed to the standard *E. coli* diet starting in generation F15. (**D)** Mean probability of the predatory mouth form on a *Novosphingobium* diet (orange) and *E. coli* (blue). The *y* axis indicates the observed Eu mouth-form frequency and the *x* axis shows the number of generations. (**E)** Mean probability of the predatory mouth form on *E. coli* after 15 and 25 generation on *Novosphingobium*, respectively. (**F)** Mean probability of the predatory mouth form on *E. coli* after 3, 5 and 10 generation on *Novosphingobium*, respectively. For E and F: The 95% HDI was calculated using a Bayesian hierarchical model. The yellow area displays the lower and upper limits of the baseline Eu response averaged across the 101 generations on *E. coli* in the control experiment.

Building on the molecular understanding of mouth-form plasticity regulation in *P. pacificus* and the vast array of tools for experimental manipulation, a recent study introduced long-term environmental induction through distinct diets and associated food-reversal experiments (Figure 1C) (Quiobe et al., 2025a). This study identified dietary induction and subsequent TEI of the predatory mouth form after food reversal (Figure 1D,E). Subsequent work established a role of the ubiquitin ligase EBAX-1/ZSWIM8 and the heat shock protein Hsp90 in TEI of the predatory mouth-form by destabilizing clustered microRNAs (Quiobe et al., 2025a). Further studies revealed that bacterial-derived vitamin B12 is necessary and sufficient to induce the TEI of the predatory mouth form (Quiobe et al., 2025b). While this work begins to elucidate the molecular machinery associated with the TEI of the predatory morph, the potential adaptive value of this phenomenon has not been investigated. Here, we test the potential adaptive value of TEI of the predatory mouth-form. We use brood size as proxy for fitness and compare naïve, diet-induced and diet-reverted animals. This study provides strong evidence for the adaptive significance of the TEI of the predatory morph in *P. pacificus*.

## Materials and Methods

### Bacterial and nematode culture maintenance

The ancestral *P. pacificus* RSC011 isolate was cryopreserved within the first 10 generations after its initial collection from Coteau Kerveguen, La Réunion. Worms were reared at 20 °C under standard conditions on Nematode Growth Medium (NGM) plates seeded with either *E. coli* OP50 or *Novosphingobium* L76. Bacterial strains were cultured overnight in Lysogeny Broth (LB) with kanamycin (50 μg/ml) when required. Specifically, OP50 was grown overnight at 37°C in LB without shaking while *Novosphingobium* L76 was grown at 30 °C at 180 rpm. NGM plates were seeded with 300 μl of overnight cultures and incubated for 2 days prior to use.

### Dietary induction and food reversals

Dietary induction on *Novosphingobium* and subsequent reversal experiments on *E. coli* OP50 were based on the long-term environmental induction (LTEI) protocol described previously (Quiobe et al., 2025a). For this study, worms were propagated on *Novosphingobium* for 5 and 10 generations, respectively. For food reversals after the F5 and F10 generations, worms were first transferred from *Novosphingobium* L76 to NGM plates supplemented with 50 μg/ml of kanamycin with a kanamycin resistant *E. coli* strain as food for one generation, before being reintroduced to standard *E. coli* OP50-seeded NGM plates for subsequent generations. Reversal experiments were carried out for 3-4 generations following the initial *Novosphingobium* exposure.

### Assays for lifetime reproductive success

To obtain synchronized worm cultures, a bleaching process is widely used in protocols for assessing brood size. However, exposure to stressors such as bleaching, starvation, or extreme temperature fluctuation are known to directly alter the mouth-form phenotype. Therefore, to avoid effects that could directly or indirectly influence life-history phenotypes, no bleaching was performed. To assess lifetime reproductive success, J4 animals were individually transferred on separate plates with 75µl of bacterial lawn (Figure 2A). Upon reaching adulthood, individual worms were transferred daily to fresh plates for four days. Almost 90% of self-progeny are laid within the first 3 days. Worms were allowed to lay eggs on the day-4 plates for several additional days to provide an accurate estimate for the remaining ∼10% of self-progeny. For lifetime reproductive success (LRS) assays, 50-60 independent replicates (mothers) were measured for each generation.

**Figure 2.**
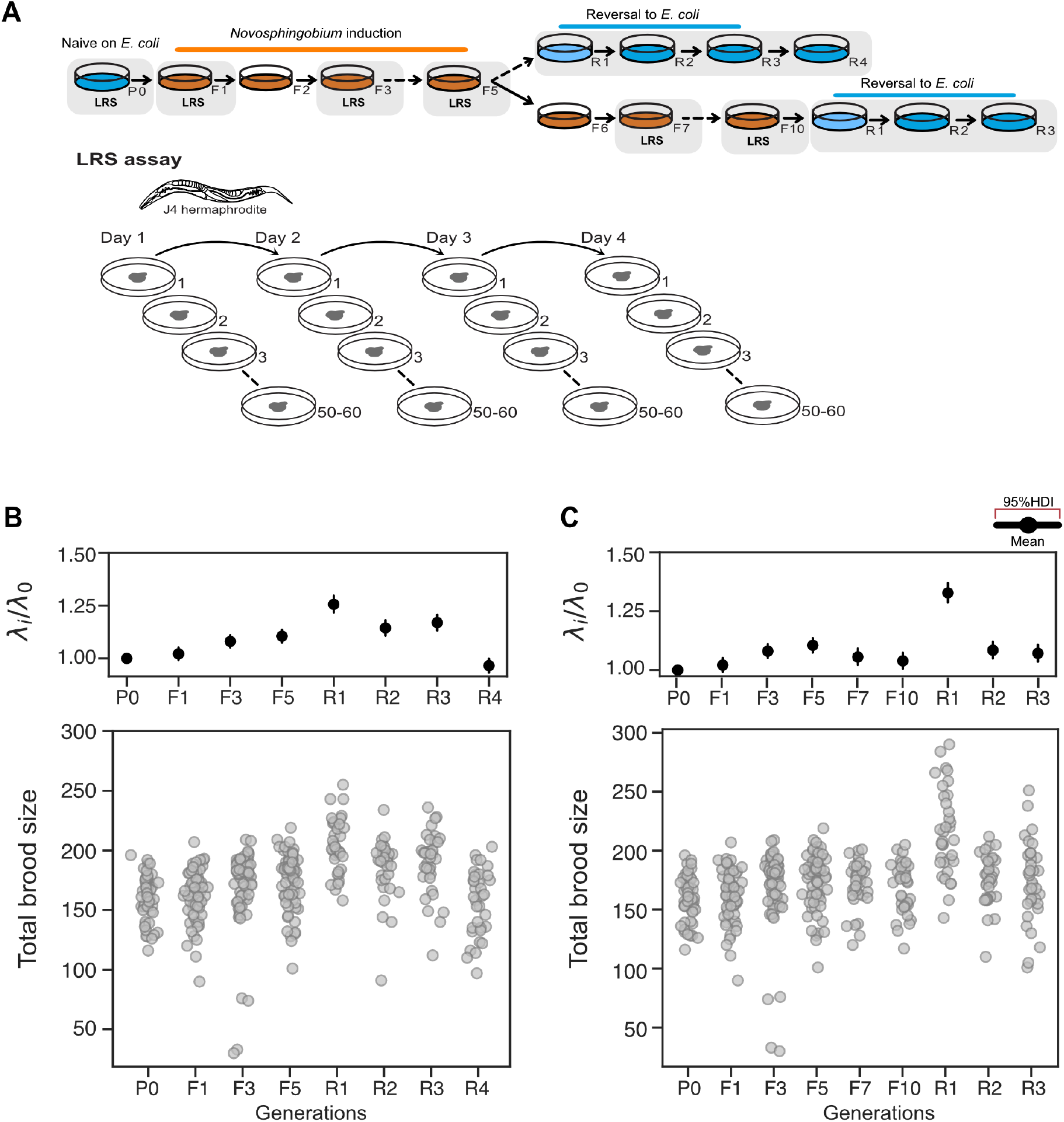
Assessment of lifetime reproductive success in environmental induction experiments. **(A)** Schematic representation of the experimental setup. Upper region: The assay for lifetime reproductive success (LRS) was performed in generations indicated in grey. Lower region: LRS was performed by transferring J4 hermaphrodites to fresh plates every 24 hours for 4 days. **(B)** Overall fecundity as a representation of LRS during the F5 reversal experiment. P0, naïve *P. pacificus* on the *E. coli* diet before the transfer to *Novosphingobium*. F1, F3 and F5 generations on *Novosphingobium* during induction. R1-R4, after reversal back to *E. coli*. Each dot represents an independent replicate. **(C)** Lifetime reproductive success during the F10 generation reversal experiment. P0, naïve *P. pacificus* on the *E. coli* diet before the transfer to *Novosphingobium*. F1, F3, F5, F7 and F10 generations during induction on *Novosphingobium*. R1-R3, after reversal back to *E. coli*. Each dot represents an independent replicate. The top panel for each (B) and (C) shows the 95% highest density interval (HDI) of the estimated ratio 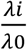. *λ*0, P0 generation; *i*, for any given generation.

### Statistical analysis

To estimate the rate parameter *λ* of the LRS, we implemented a Bayesian model using PyMC version 5.16.2 (Abril-Pla et al., 2023). For each dataset, observations were assumed as Poisson-distributed counts with the rate parameter *λ*. We assume the rate *λ* to have a Γ(*α* = 1, *β* = 2) prior distribution. Posterior inference was performed using No-U-Turn Sampler (NUTS) by running 3,000 tuning iterations followed by 5,000 sampling iterations. Next, the distribution ratio 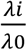 was obtained, where *λ*0 corresponds to the inferred distribution for *λ* given P0 and the corresponding distribution for treatment *i*. The 95% highest density interval (HDI) of this ratio was then used to compare LRS across generations.

## Results and Discussion

### Background: Long-term environmental induction experiments in *P. pacificus*

A long-term environmental induction set up was recently established for mouth-form developmental plasticity in *P. pacificus* (Quiobe et al., 2025a). Specifically, the natural isolate *P. pacificus* RSC011 from a high-altitude location on La Réunion Island, which has a mouth-form ratio of 30%:70% Eu:St worms when cultured on the standard laboratory food *E. coli* OP50, was used for the experiment (Figure 1C). Starting from the total offspring of a single hermaphrodite, 110 genetically identical lines were exposed to a *Novosphingobium* diet, a bacterium derived from the natural environment of *Pristionchus* (Akduman et al., 2020). Worm lines were propagated for 101 generations by single worm descent and starting in generation F15, all lines were reverted to the ancestral *E. coli* diet all 10 generations (i.e. F25, F35 etc). We found dietary induction of the predatory mouth form: all lines showed an immediate and systemic response to the new diet by expressing the Eu mouth form in all animals and throughout the entire experiment (Figure 1D). After reversal to *E. coli*, we observed high probability of developing the Eu mouth form for more than three generations before the mouth-form ratio dropped to the baseline level of naïve animals (around 30% Eu); thus, representing an example of transgenerational inheritance (Figure 1E). Subsequent experiments indicated that a minimal exposure to *Novosphingobium* of five generations is necessary to induce this TEI of the predatory morph. In contrast, return to the *E. coli* diet after one, two or three generations on *Novosphingobium*, does not result in any memory (Quiobe et al., 2025a) (Figure 1F). Therefore, standard assays to test for TEI of the predatory mouth form involve 5 and 10 generation exposures to *Novosphingobium* before reversal to *E. coli*. This example of diet-induced expression of the predatory mouth-form is a striking example of the inheritance of non-genetic information and the associated molecular machinery in both, the bacterial diet and the responding worms are being investigated (Quiobe et al., 2025a; Quiobe et al., 2025b). However, whether the induction and subsequent memory of the predatory morph is correlated with increased fitness has not been investigated. Thus, the ecologically relevance and the potential adaptive value of this example of multigenerational plasticity have not been determined, requiring further investigations.

### Criteria for the adaptive value of transgenerational inheritance

In a recent review article, Baugh and Day have proposed that three criteria have to be fulfilled to document the adaptive value of transgenerational inheritance and multigenerational plasticity (Baugh & Day, 2020). First, it has to be demonstrated that an environmental response of the organism is transmitted across generations. In outbreeding species this can be challenging because selection of existing genetic variation has to be ruled out. However, in the self-fertilizing *P. pacificus*, the dietary induction of the predatory mouth form described above fulfills this criterion because there is no standing genetic variation in these cultures (Figure 1) (Quiobe, et al., 2025a). Second, an explanation for the mechanistic basis of the phenomenon must be provided (Baugh & Day, 2020). This requires the identification of the molecular and cellular mechanisms involved in the heritable transmission of non-genetic information. Such mechanistic insight is necessary to rule out the possibility that spontaneous mutations or other confounding factors are responsible for the observed phenomenon. In the case of *P. pacificus* mouth-form memory, genetic and experimental approaches have identified vitamin B12 as the inducing agent from the *Novosphingobium* diet (Quiobe et al., 2025b) and a role of the ubiquitin ligase EBAX-1/ZSWIM8 through destabilization of clustered miRNAs in the worm as crucial factors for memory formation (Quiobe et al., 2025a). Thus, *P. pacificus* transgenerational inheritance of the predatory mouth-form fulfills the first two criteria proposed by Baugh and Day. The third criterion is the crucial demonstration that multigenerational effects beyond the original exposure to an environmental stimulus improves descendant fitness (Baugh & Day, 2020). Below, we describe experiments using brood size as proxy for fitness to study the adaptive value of TEI of the predatory morph in *P. pacificus* RSC011 and also study developmental speed as another life history trait.

### Assessing fitness aspects in environmental induction experiments

In the nematode androdioecious hermaphrodites *C. elegans* and *P. pacificus*, total brood size after self-fertilization represents the most commonly used proxy for fitness (Dardiry et al., 2023; Pete & Hunter, 2025). As the total offspring of a hermaphrodite is nearly exclusively hermaphroditic and can therefore reproduce itself, brood size is directly related to the reproductive value as originally defined by Fisher (Fisher, 1930). In such hermaphrodites, this measurement is equivalent to the lifetime reproductive success (LRS), which represents the total number of viable offspring an individual produces throughout its entire life. In this study, we use the term ‘LRS’ to evaluate the potential adaptive value of transgenerational inheritance of the predatory mouth form. Specifically, we measured LRS of single *P. pacificus* RSC011 hermaphrodites in three different conditions; i) during naive *E. coli* OP50 conditions, ii) during *Novosphingobium* exposure for 5 or 10 generations, respectively; and iii) after food reversal back to *E. coli* (Figure 2A). For each condition, LRS measurements started with J4 animals transferred to nematode growth medium (NGM) agar plates seeded with either *E. coli* or *Novosphingobium*. Testing for daily fecundity showed that the majority of progenies were laid within a window of four days upon reaching adulthood (Figure 2A).

We found that animals grown on the standard *E. coli* OP50 food produced ∼ 160 adult offspring. Upon switching RSC011 worms to *Novosphingobium* for one generation, LRS remained similar to naïve populations grown on *E. coli* OP50 (Figure 2B). A significant increase in LRS to an average of 170 and 175 offspring was observed in the F3 and F5 generations on *Novosphingobium*, respectively (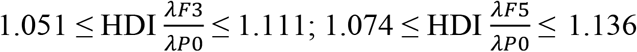) (Figure 2B). This result highlights that the LRS increased on *Novosphingobium*, however, not immediately as it was previously seen for the dietary induction of the predatory Eu mouth form (Figure 1D) (Quiobe, et al., 2025a). In addition, both F7 and F10 generations on *Novosphingobium* had a slight decrease in LRS when compared to the F3 and F5 generations (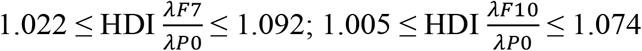) (Figure 2C). However, the LRS of the F7 and F10 generations were still significantly higher than those of P0 animals (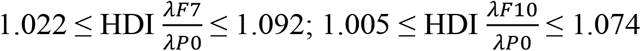) (Figure 2C). Taken together, feeding *P. pacificus* RSC011 on *Novosphingobium* resulted in an increased LRS relative to *E. coli* grown worms indicating that the change of mouth form on this diet is accompanied by a fitness advantage.

### Reversal from a *Novosphingobium* diet strongly increases brood size in *P. pacificus*

To assess whether the higher LRS observed during exposure to *Novosphingobium* can be inherited, we transferred worms from *Novosphingobium* to NGM plates supplemented with kanamycin and used a kanamycin-resistant *E. coli* strain as food source for one generation. Kanamycin-supplemented plates with kanamycin-resistant *E. coli* on their own had no significant effect on LRS when standard *E. coli* OP50-grown worms were transferred to kanamycin-resistant *E. coli* (Supplementary Figure 1). In contrast, we observed a significant increase in LRS relative to naïve P0 animals upon reversal back to *E. coli*. Specifically, F5R1 animals following the *Novosphingobium* exposure for five generations on average produced 204 offspring (Figure 2B,C). This finding indicates a significant increase in LRS compared to P0 animals and to all generations on *Novopshingobium* (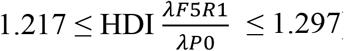). Furthermore, an increase of the LRS was also observed in the F5R2 and F5R3 generations (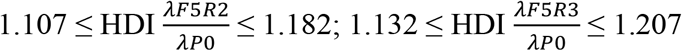) (Figure 2B). Similarly, after the *Novosphingobium* exposure for 10 generations, an increase of the LRS relative to P0 animals was seen in the F10R1, F10R2 and F10R3 generations (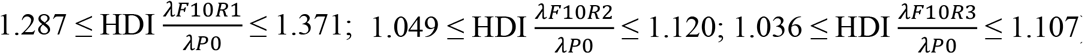) (Figure 2C). Note that in both reversal experiments, the F5R1 and F10R1 generations showed the highest increase in total LRS. These results indicate that the increase in LRS can be inherited for up to three generations after a multigenerational exposure to the *Novosphingobium* diet.

### An adaptive value of the transgenerational inheritance of the predatory mouth form

Transgenerational inheritance is widely observed in organisms, yet its adaptive value remains controversial (Santilli & Boskovic, 2023). Rather than persisting indefinitely, TEI follows a tunable duration suggesting that inheritance duration is evolutionary optimized (Houri-Ze’evi et al., 2016). Our study aimed to investigate the total brood size or LRS as a well-established proxy for fitness in the context of food reversal experiments in *P. pacificus*. Our results show that exposure of *P. pacificus* RSC011 worms to *Novosphingobium* increases LRS and is inherited transgenerationally. Our observations add to the response patterns of nematodes to dietary switch experiments. Exposure of *C. elegans* to a vitamin B12-rich *Comamonas* diet results in faster development but with reduced LRS (MacNeil et al., 2013; Watson et al., 2014). In contrast, two *P. pacificus* wild isolates exhibited different outcomes depending on their mouth-form preferences; one strain had an increased, the other one a decreased LRS (Dardiry et al., 2023). However, these studies did not test for LRS over multiple generations and they did not involve food reversal experiments. Our findings highlight the strain-specific effects, indicating that after the original dietary induction of the predatory mouth form, brood size is higher for multiple generations when compared to naïve animals. Both, the Eu morph and LRS are transgenerationally inherited during food reversal for multiple generations suggesting that the transgenerational memory of the predatory mouth form has an adaptive value. To our knowledge this study is the first to observe such effects in the context of food reversal and transgenerational inheritance.

The previous study by Dardiry and co-workers mentioned above had used dietary switch experiments to investigate the potential ‘cost of plasticity’ in *P. pacificus* (Dardiry et al., 2023). Studying several *P. pacificus* wild isolates revealed that in response to a *Novosphingobium* diet, worms produced a higher proportion of the Eu morph. However, the increase in the Eu mouth form was accompanied by the production of fewer offspring illustrating a cost of plasticity (Dardiry et al., 2023). It is important to note that these experiments have measured life history traits in the first generation after the dietary switch. In contrast, we have followed reproductive fitness for 5 and 10 generations. Therefore, we suggest that future studies using similar experimental approaches should investigate life history traits over multiple generations. Overall, our study shows that after dietary switch and food reversal *P. pacificus* RSC011 has a higher LRS for multiple generations suggesting that the expression of the predatory mouth form has an adaptive value.

Finally, it is important to note that transgenerational inheritance has to be considered in the context of bet-hedging strategies. Environmental changes in space and time are challenging for organisms and can be broadly categorized into environmental heterogeneity and fluctuating (unpredictable) environments. The scarab beetle carcasses on which *P. pacificus* are reliably found (Herrmann, Mayer & Sommer, 2006; Kanzaki et al., 2021), represent fluctuating environments where the duration of food availability cannot easily be predicted (Renahan & Sommer, 2021; Renahan et al., 2021). Many large animals can respond to such environmental changes through matching habitat choice (Edelaar, Siepielski & Clobert, 2008), whereas bacteria, plants but also soil-dwelling nematodes cannot. In bacteria, stochastic phenotypic switching is a known bet-hedging strategy to overcome environmental change (Rainey et al., 2011). In nematodes, transgenerational inheritance might be considered another bet-hedging strategy allowing offspring to be pre-adapted to environmental conditions experienced by previous generations. Several recent studies in *C. elegans* discuss such bet-hedging strategies. For example, starvation-induced small RNAs transmitted to progeny can extend lifespan and alter metabolism, enhancing survival under recurring food scarcity, but potentially imposing costs when resources are plentiful (Rechavi et al., 2014). Similarly, temporary dietary restriction in parents improves late-life reproduction, yet later-generation descendants exhibit reduced fitness under normal feeding conditions (Ivimey-Cook et al., 2021). Behavioral adaptations, such as learned avoidance of pathogens (Moore, Kaletsky & Murphy, 2019) or stress-induced pheromone signaling that promotes outcrossing (Toker et al., 2022), also prepare offspring for potentially recurring threats while introducing trade-offs in less stressful environments. In *P. pacificus* RSC011 worms, the increased Eu mouth form and higher LRS and their transgenerational inheritance confer a fitness advantage. This likely supports adaptation of *P. pacificus* to patchy, ephemeral environments that it might encounter in the wild. Furthermore, the tunable duration of the mouth-form memory suggests that worms can calibrate the persistence of such responses by balancing the benefits of anticipatory adaptation to recurring environments against the risks of maladaptation. Taken together, these findings support the view that TEI in nematodes enable them to hedge their bets by spreading the consequences of an environmentally-induced decision across generations.

## Acknowledgments

This work was funded by the Max-Planck Society. We thank Dr. Ata Kalirad for fruitful discussions and help with the graphical representation. We would like to thank the entire Sommer lab for continuous support throughout this project.

## Author contributions

Shiela Pearl Quiobe (Conceptualization-Equal, Data curation-Lead, Formal analysis-Lead, Investigation-Lead, Methodology-Lead, Visualization-Equal, Writing - original draft-Equal)

Ralf Sommer (Conceptualization-Equal, Funding acquisition-Lead, Investigation-Equal, Resources-Supporting, Supervision-Lead, Writing - original draft-Lead, Writing - review & editing-Lead)

## Conflict of interest

The authors declare no competing interests.

## Data availability statement

The data is available at https://github.com/shielapearl18/Adaptive-consequences-of-TEI-of-the-predatory-morph

## Notes

### Competing Interest Statement

The authors have declared no competing interest.

