## Supplemental Figure 1 for "Adaptive consequences of transgenerational inheritance of a predatory mouth-form trait in nematodes"

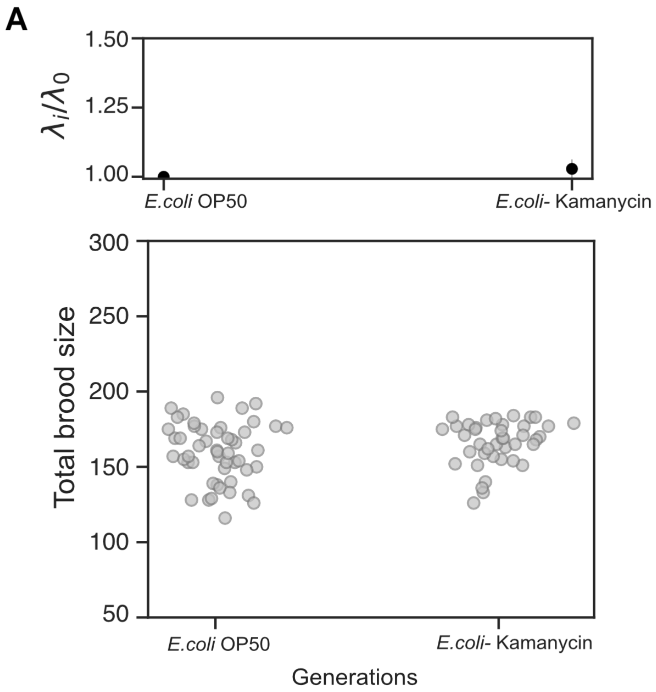


**Supplementary Figure 1. Assessment of the lifetime reproductive success of animals on kanamycin- supplemented plates.**

The top panel shows the 95% highest density interval (HDI) of the estimated ratio $\frac{\lambda i}{\lambda 0}$ . $\lambda 0$, P0 generation; $i$, for the kanamycin-exposed generation.
